# The GRE Over the Entire Range of Scores Lacks Predictive Ability for PhD Outcomes In the Biomedical Sciences

**DOI:** 10.1101/373225

**Authors:** Linda Sealy, Christina Saunders, Jeffrey Blume, Roger Chalkley

## Abstract

The association between GRE scores and academic success in graduate programs is currently of national interest. GRE scores are often assumed to be predictive of student success in graduate school. However, we found no such association in admission data from Vanderbilt’s Initiative for Maximizing Student Diversity (IMSD), which recruited historically underrepresented students for graduate study in the biomedical sciences at Vanderbilt University spanning a wide range of GRE scores. This study avoids the typical biases of most GRE investigations of performance where only high-achievers on the GRE were admitted. GRE scores, while collected at admission, were not used or consulted for admission decisions and comprise the full range of percentiles, from 1% to 91%. We report on the 29 students recruited to the Vanderbilt IMSD from 2007-2011 who have completed the program at this date. While the data set is not large, the predictive trends between GRE and long-term graduate outcomes (publications, first author publications, time to degree, predoctoral fellowship awards, and faculty evaluations) are remarkably null and there is sufficient precision to rule out even mild relationships between GRE and these outcomes. Career outcomes are encouraging; many students are in postdocs, and the rest are in stage-appropriate career environments for such a cohort, including tenure track faculty, biotech and entrepreneurship careers.

## Introduction

Recently Moneta-Kohler et al. [1] published a detailed statistical analysis of the lack of ability of the GRE to predict performance in graduate school in the biomedical research arena at Vanderbilt. A similar study was published by Hall et al. [2] from the University of North Carolina Chapel Hill. However, there was a limitation to the overall conclusions in that the range of GRE scores did not cover scores lower than approximately 50%. In order to test if such a limitation impacted the predictive ability of the GRE, we would need to admit students for whom we had GRE information, but where the admitted students covered the entire range of scores with no bias or cut-off (deliberate or otherwise) in the level of the score. This is a difficult requirement, as admissions committees normally do not pursue applicants with very low GRE scores, even if other aspects of the application might appear to be competitive. We are aware that a fairly significant number of schools are electing to not use GRE scores at all in making admissions decisions [3]. Other schools may be considering whether or not to require GRE scores, but have not yet taken action. All of these schools surely might benefit if there were to be an experiment in which we assayed the predictive ability of the GRE scores over the entire range of scores.

We report that we have performed this natural experiment with GRE scores covering the range from 1st to 91st percentile, in an approach where the scores, although submitted as part of the application, were not considered in the selection of incoming graduate students. This came about in the following way. In 2007 Vanderbilt was awarded an NIGMS-funded IMSD program with the goal of increasing the number of students from underrepresented groups completing PhDs in the biomedical sciences. This program was a redesign of our previous IMSD post baccalaureate program in response to the NIH stipulation in 2006 that students in the program had to be matriculated as graduate students, not post baccalaureates. We were aware that increasing the number of historically underrepresented (UR) students in our PhD programs might result in another school(s) not enrolling these students, and the overall pool of UR PhD trainees would remain static. This was because at that time the pool of high qualified UR students, when quantitative metrics (GRE, GPA) were a key driver of the assessment, was in insufficient supply. The authors had already collected data over a ten-year period indicating that for underrepresented students at least the GRE at the levels usually expected for admission offered no guidance in terms of achievements of long term PhD training goals. Consequently, we decided that removing the barrier of GRE scores to admission would actually lead to an overall increase in historically underrepresented PhD trainees.

Therefore, in 2007 the Vanderbilt IMSD program adopted a fully holistic approach to admissions. The GRE scores were recorded as a required part of the application process, but they were essentially ignored by the IMSD admissions committee, which operated in a separate fashion from our regular interdisciplinary graduate program (IGP) admission committee. This resulted in a group of students who were eligible for IMSD support (as defined by NIGMS) admitted with GRE scores over the full range (1-90% GRE-V and 11-91% GRE-Q). Details of the fully holistic approach are presented below, but relied heavily on letters of recommendation, personal statements and interviews. If these factors were strong, no GRE score was too low to be admitted.

Over the next four years, 29 students,were admitted in this fashion, all of whom have now completed the PhD graduation cycle, so we are able to evaluate the outcomes of admissions strategies which cover the entire GRE range (including both very high and low scores), under conditions in which the admissions process operated obliviously to the scores themselves. The measures we have used to evaluate outcomes performance in biomedical research were also used in our previous report [1] on the lack of predictive value of the GRE. These include: number of first or other order author papers, receiving competitive fellowship awards, time to degree, a detailed faculty evaluation at the time of graduation, and an initial review of career development in scientific areas. Of the 29 URM students who participated in the IMSD program over this time period, 27 have now graduated (25 PhD, 2 MS) with two students dropping out early as a consequence of health problems. We present here the outcomes of the 25 PhD graduates and the relationship of these outcomes to their GRE scores.

## Materials and Methods

GRE (Quantitative and Verbal) scores and academic performance data from 25 IMSD students who matriculated from 2007 to 2011 were collected and examined. Academic performance outcomes of interest were: time elapsed in program (i.e., months to degree), number of publications, number of first-author publications, fellowship status (any or F31), Vanderbilt faculty ranking (10 = best, 50 = worst). Table 2 provides univariate summaries (e.g., mean, median, standard deviation, inter-quartile range) of these variables. Figure 2 presents histograms of the continuous outcome variables. Regression modeling was used to assess the degree of association between GRE outcomes and academic outcomes. Specifically, Poisson regression was used to model publication counts (accounting for length of time in the program), months to degree, and faculty ranking. Logistic regression was used to model receipt of fellowship. For all models, we report point estimates, model robust standard errors, and 95% confidence intervals (CIs). We plot each performance measure as a function of GRE scores and include the fitted regression line as well as a locally weighted scatterplot smoother (lowess) line to visually assess linearity assumptions and model fit. Confidence intervals were plotted to demonstrate the degree of precision afforded by the data at the 95% level. Any relationship between GRE scores and outcomes would be captured in the slope of these regression lines. While it is not possible to prove the null hypothesis that GRE scores and outcomes are not related, it is possible to provide an upper bound on the largest potential association. The 95% CI provide this bound and comprise the set of associations supported by the data. As we will see from the data, despite the small sample size, these CIs do not support mild or strong associations between GRE scores and outcomes. For a sensitivity analysis, we compared academic outcomes between the first quartile and the fourth quartile of GRE scores. If any association were present, such an analysis should at least yield exaggerated point estimates of the association effect. The research was approved by Vanderbilt University IRB (151678). Consent was not given as data were analyzed anonymously.

## Results

In Fig 1 we report the range of GRE scores among the 27 URM students admitted into the graduate program in the biomedical sciences at Vanderbilt from 2007 through 2011 and who completed a PhD or MS degree. The admission decisions for these students during this time period was determined by the IMSD admissions/advisory committee, and although the student’s GRE score was recorded in our databases, it has only been used for outcomes studies long after the admissions event. The range of GRE scores among this group of students varies across the spectrum for students who were admitted in response to a detailed analysis of the committee’s assessment of the likelihood of the student’s success in research. The committee’s assessment was based primarily on the non-quantitative components of the application, including a close reading of the letters of recommendation and the student’s personal statement. The student’s transcript was evaluated, primarily to assess adequate coursework preparation for biomedical PhD coursework. A wide range of GPAs were accepted. We sought to place the overall and science GPAs in the context of the college or university and the life events of the applicant. For example, students with extensive work and/or family responsibilities might reasonably be expected to end up with lower GPAs due to time demands. The lowest GPA accepted among this group of students was 1.8. Finally, all students were invited to campus for an interview visit that was also given significant consideration.

**Fig 1.**
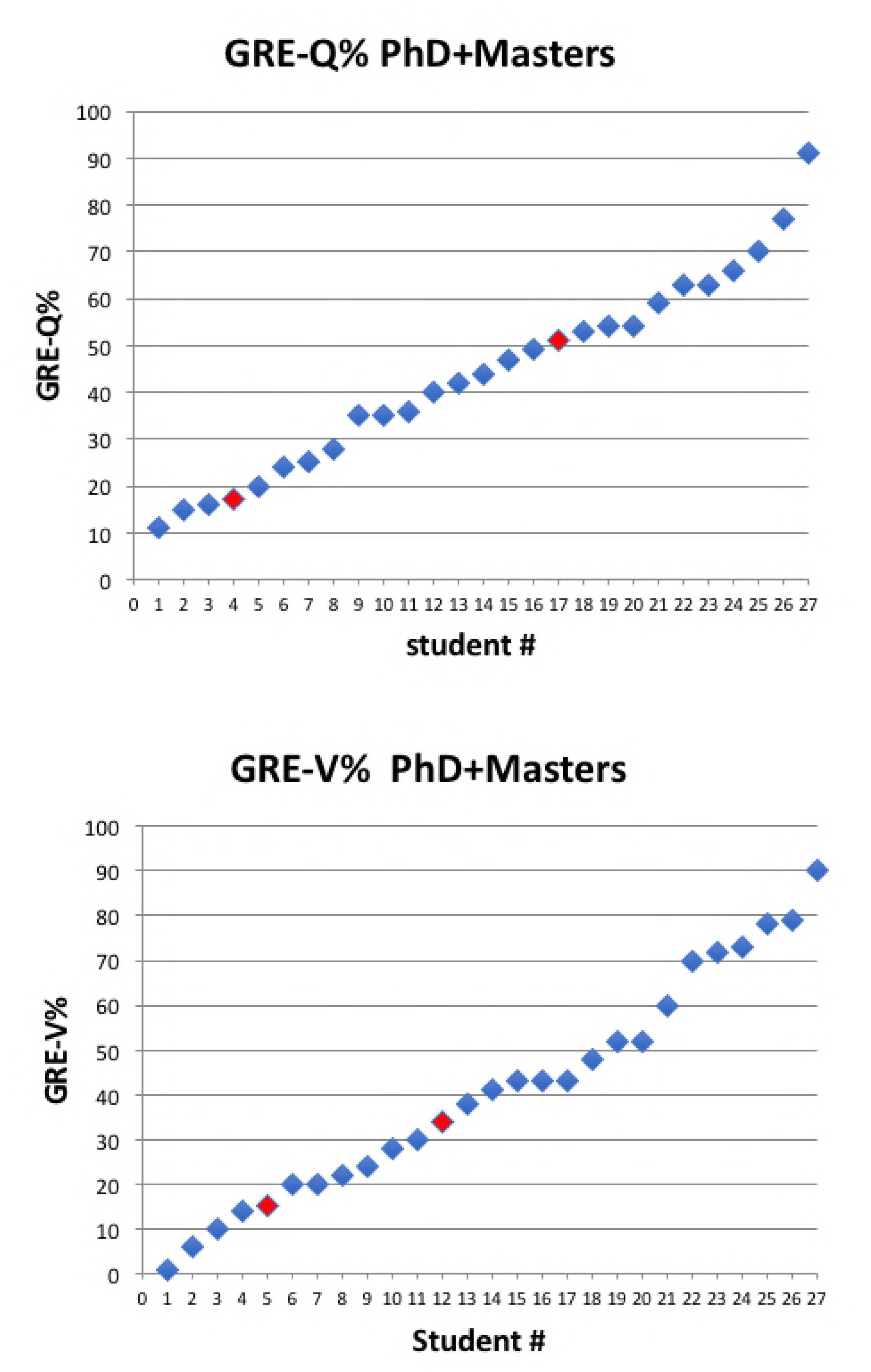
GRE Quantitative and Verbal Scores of IMSD Students Matriculating from 2007-11 who completed a PhD or Master’s degree. Top panel depicts GRE-Q% and lower panel depicts GRE-V% for 27 students who completed either a PhD (blue symbols) or Master’s degree (red symbols).

From the 27 students with GRE scores shown in Fig 1, 93% have graduated with the PhD and 7% left with an MS degree. Of the 25 students who completed the PhD, 84% continued to postdoctoral positions. Four students did not continue on to postdocs, choosing instead to move to industry, consulting, medical school, or an academic faculty position. Overall, the outcomes of this cadre of GRE-blind admitted students are strikingly parallel to those of students admitted through the traditional route (using much higher GRE scores) over the same time period [4]. As indicated in Fig 1, we have a wide range of GRE scores among this group. This unusual group provided us with a means to test the predictive value of GRE scores over a much wider range than most admissions committees will typically tolerate.

Fig 2 shows histograms of the data analyzed in this study: range and frequencies of GRE scores, number of publications, number of first author publications, months to degree, and faculty ranking. The faculty ranking is obtained upon the student’s completion of their Ph.D. The ranking is comprised of the sum of scores for each of ten questions, listed in Table 1. The questions cover a range of areas that are often informally assessed as measures of developing into a successful, independent scientist; many would fall into the area of the social/emotional learning skillset. We ask the PhD faculty mentor to score their newly-minted PhD student from one to five, with one being best. Thus, the top ranking possible is a 10, if the student received a score of one for each of the ten questions. Student rankings ranged from 12 to 39 with a median of 21.5. The other metrics are self-explanatory, with number of publications ranging from one to thirteen (median= 4) and first author publications from one to six (median=2). Note that students are expected to publish at least one first author paper as a requirement for the PhD in most of our PhD granting programs. The time to degree for these students ranged from slightly more than 4 years, to just over 7 years (median = 5.66 years). In addition to the data shown in Fig 2, we also included whether or not the student obtained an individual fellowship in a national competition (F31, AHA, DOD, etc) as an additional metric (see Figs 6 and 7). Summary statistics of the data for this study are presented in Table 2. The hypothesis we test is that GRE scores are associated with future performance in a biomedical graduate program. This association will be measured by the slope in a regression model, to be explained shortly.

**Fig 2.**
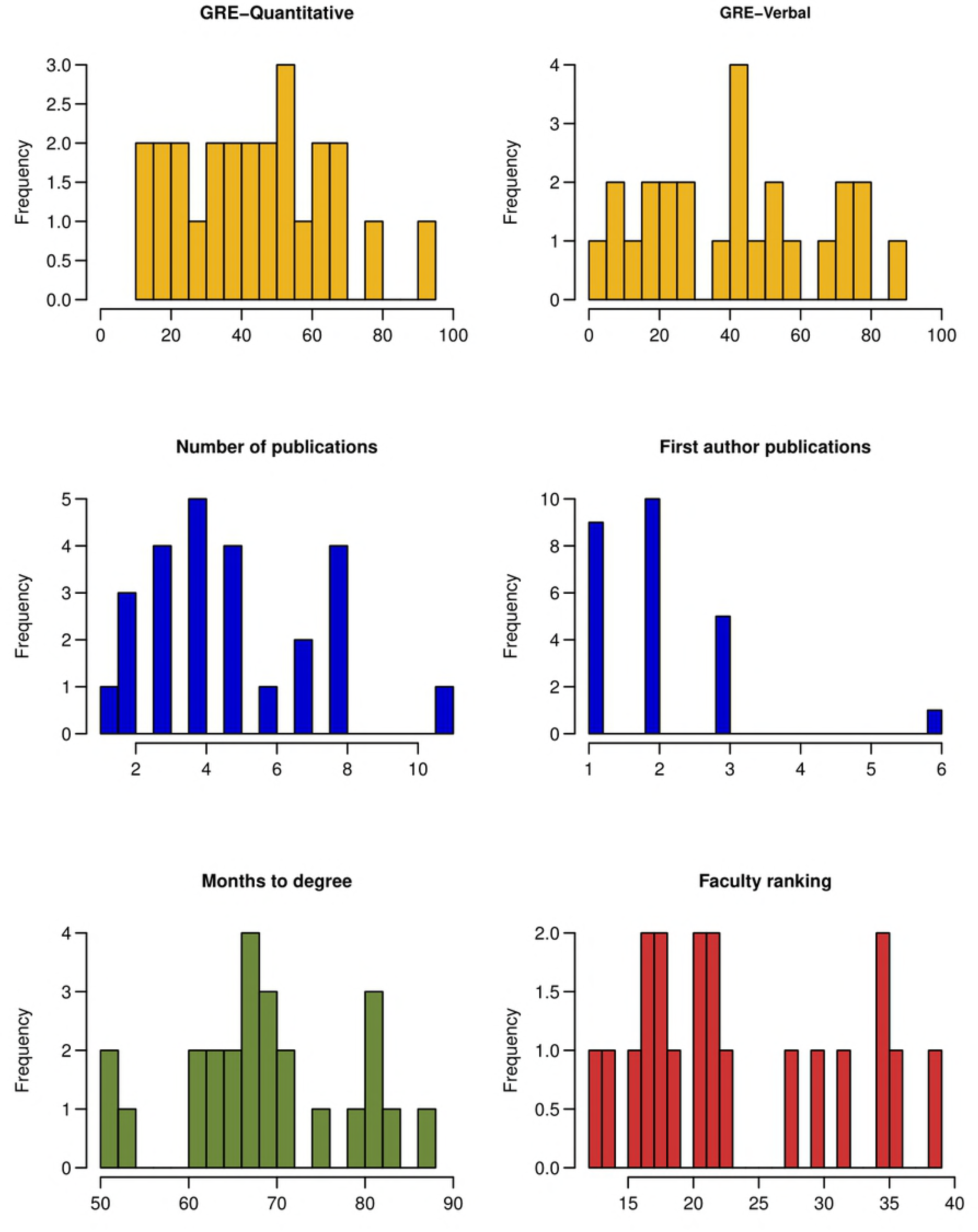
Histograms of outcomes data. Frequencies of GRE scores (Q% and V%), number of publications, months to degree, and faculty ranking are shown as indicated.

**Table 1.** Faculty rating of student at exit.

|  |
| --- |
| 1. Ability to handle classwork needed for your PhD program |
| 2. Drive and determination |
| 3. Creativity and imagination in terms of experimental design and interpretation |
| 4. Technical ability |
| 5. Keeping up with the literature |
| 6. Output – translating observations into a presentable paper |
| 7. Ability to write creatively |
| 8. Leadership in the lab and department |
| 9. Trajectory |
| 10. Overall assessment as a productive scientist |

Faculty mentors were asked to score student upon PhD completion using a scale of 1-5 as follows: 1-outstanding; 2-excellent; 3-good; 4-fair; 5-poor. The faculty rating is the sum of the scores for 10 questions.

**Table 2.**
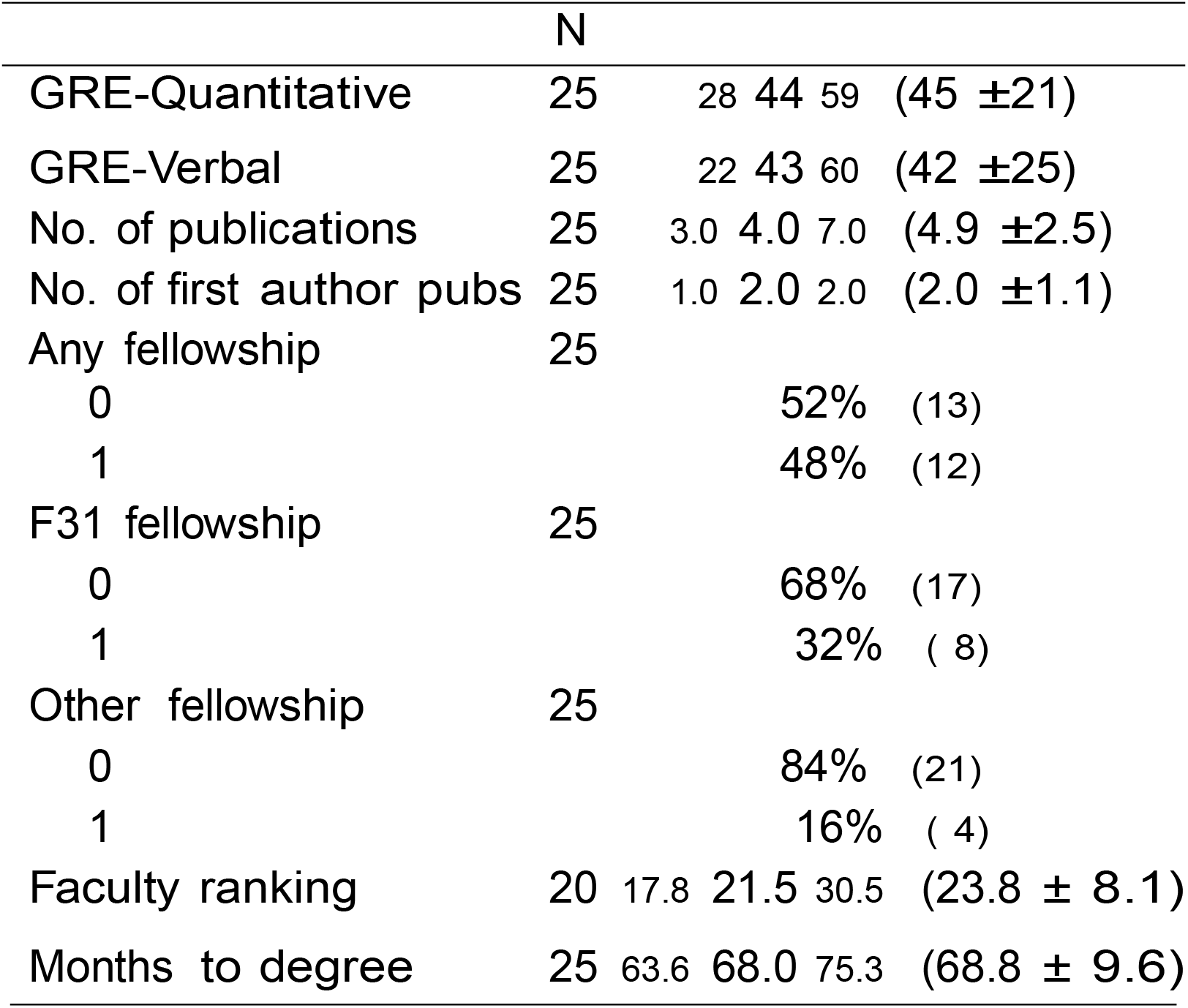
Summary statistics of GRE data.

|  | N |  |  |  |  |
| --- | --- | --- | --- | --- | --- |
| GRE-Quantitative | 25 | 28 | 44 | 59 | (45 ±21) |
| GRE-Verbal | 25 | 22 | 43 | 60 | (42 ±25) |
| No. of publications | 25 | 3.0 | 4.0 | 7.0 | (4.9 ±2.5) |
| No. of first author pubs | 25 | 1.0 | 2.0 | 2.0 | (2.0 ±1.1) |
| Any fellowship | 25 |  |  |  |  |
| 0 |  |  |  | 52% | (13) |
| 1 |  |  |  | 48% | (12) |
| F31 fellowship | 25 |  |  |  |  |
| 0 |  |  |  | 68% | (17) |
| 1 |  |  |  | 32% | ( 8) |
| Other fellowship | 25 |  |  |  |  |
| 0 |  |  |  | 84% | (21) |
| 1 |  |  |  | 16% | ( 4) |
| Faculty ranking | 20 | 17.8 | 21.5 | 30.5 | (23.8 ± 8.1) |
| Months to degree | 25 | 63.6 | 68.0 | 75.3 | (68.8 ± 9.6) |

*a b c* represent the lower quartile *a*, the median *b*, and the upper quartile *c* for continuous variables. *x* ± *s* represents *X̅* ± 1 standard deviation. *N* is the number of non−missing values. Numbers after percents are frequencies.

### Lack of association between GRE scores and publications

We modeled the relationship between total number of publications and GRE scores using Poisson regression in Fig 3 for GRE-Q (left panel) and GRE-V (right panel). Solid curves show the fitted values from the regression models (dashed lines are 95% confidence intervals) and the grey curves show lowess smoothers (locally weighted scatterplot smoother). Increasing a student’s GRE-Q score by 20 points increases their expected publication rate by just 3% (rate ratio = 1.03 with 95% CI 0.85 to 1.26). For instance, students with GRE-Q scores of 40 and 60 are expected to have 4.79 and 4.94 publications, respectively. Interesting, increasing a student’s GRE-V score by 20 points decreases their expected publication rate by 4% (0.77, 1.19). For instance, students with GRE-V scores of 40 and 60 are expected to have 4.83 and 4.62 publications, respectively.

**Fig 3.**
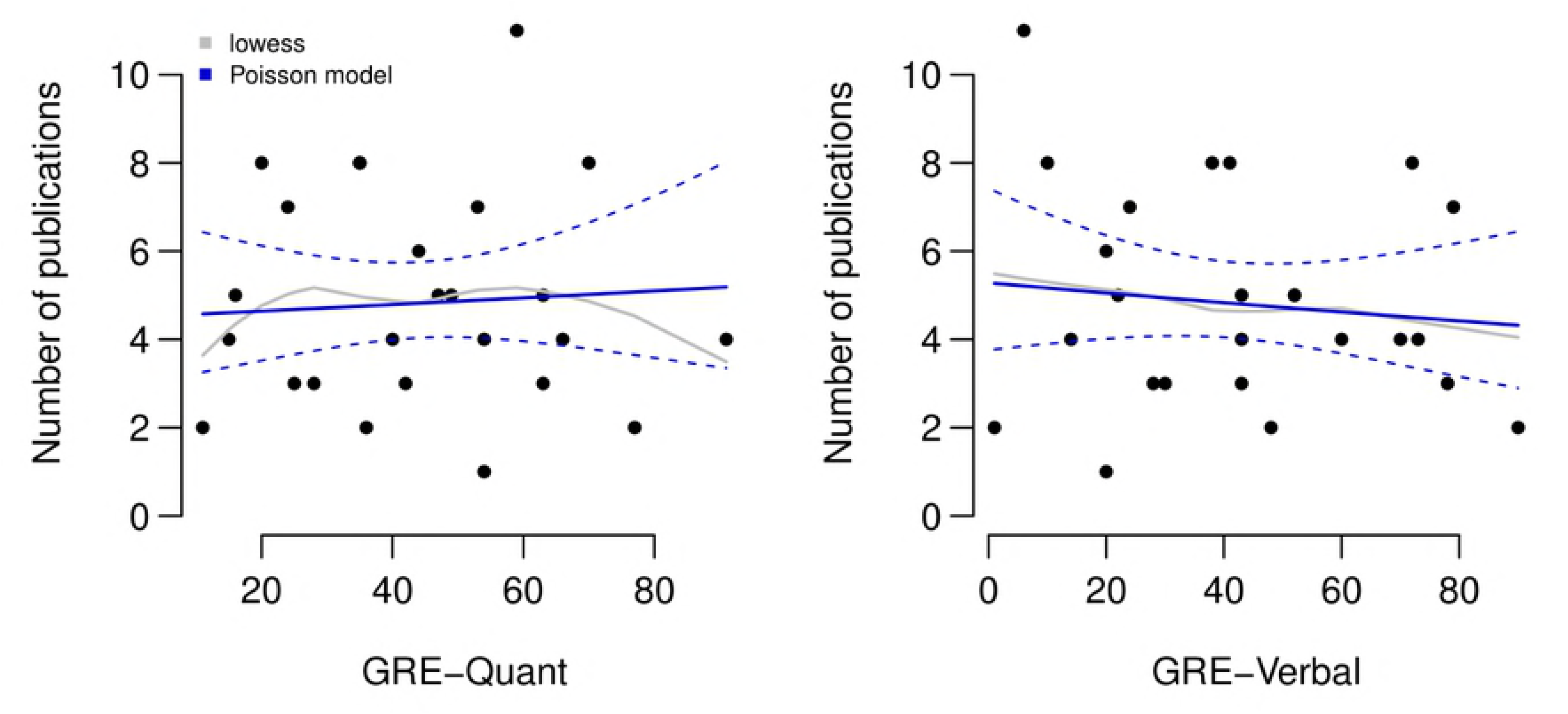
Associations between quantitative and verbal GRE scores and total number of publications. Solid curves show the fitted values from the Poisson regression models (dashed lines are 95% confidence intervals) and the grey curves show lowess smoothers (locally weighted scatterplot smoother).

We do not judge these minor differences to be significant, although the same negative correlation with GRE-V scores was also observed when the total number of first author publications and GRE scores was modeled using Poisson regression in Fig 4. Increasing a student’s GRE-V score by 20 points decreases their expected publication rate by 13% (rate ratio =0.87 with 95% CI 0.69 to 1.11). For instance, students with GRE-V scores of 40 and 60 are expected to have1.97 and 1.72 first author publications, respectively. Increasing a student’s GRE-Q score by 20 points increases their expected first author publication rate by 8% (rate ratio = 1.077 with 95% CI 0.88 to 1.32). For instance, students with GRE-Q scores of 40 and 60 are expected to have 1.94 and 2.09 first author publications, respectively, which is essentially no difference. We conclude that even when GRE scores below 20 percentile are in the mix, productivity as measured by the key currency of the scientific enterprise, namely publications - exhibits very little dependence, if any, on GRE scores.

**Fig 4.**
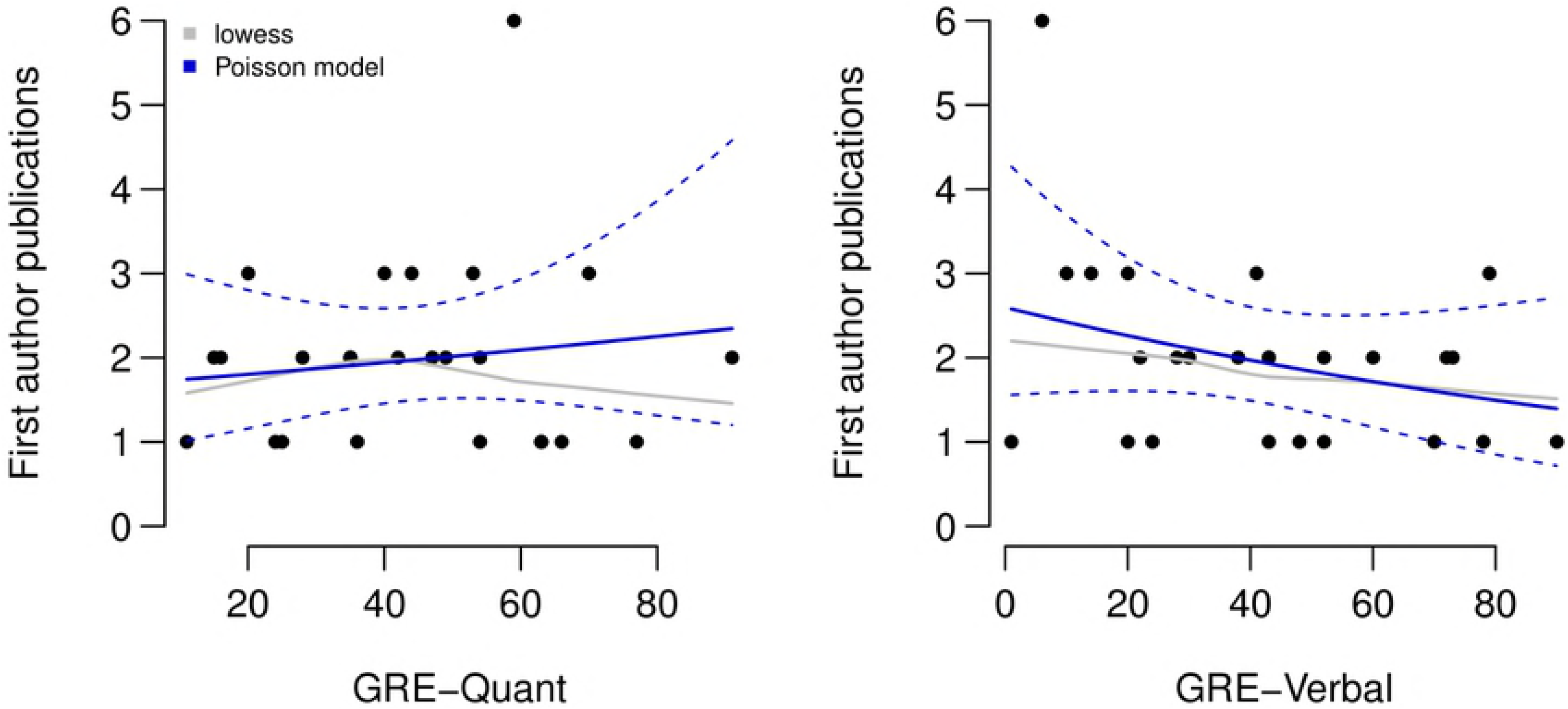
Associations between quantitative and verbal GRE scores and total number of first author publications. Solid curves show the fitted values from the Poisson regression models (dashed lines are 95% confidence intervals) and the grey curves show lowess smoothers (locally weighted scatterplot smoother).

### Lack of association between GRE scores and time to degree

In Fig 5, months to degree is plotted vs either GRE-Q (left panel) or GRE-V (right panel). Again, the solid curve shows the fitted values from the Poisson regression model (dashed lines are 95% confidence intervals) and the grey curve shows a lowess smoother (locally weighted scatterplot smoother). We observe only a very minor correlation between higher GRE scores and shorter time to degree. Increasing either the GRE-Q or GRE-V by 20 percentage points leads to a decrease in expected time to degree attainment of 3% (rate ratio = 0.97 with 95% CI 0.92 to 1.01) or 2% (rate ratio = 0.98 with 95% CI 0.93 to 1.03), respectively. This means that students with GRE-Q scores of 40 and 60 are expected to take 69 months and 67 months to complete their degree, respectively. Likewise, students with GRE-V scores of 40 and 60 are expected to take 69 months and 68 months to complete their degree, respectively.

**Fig 5.**
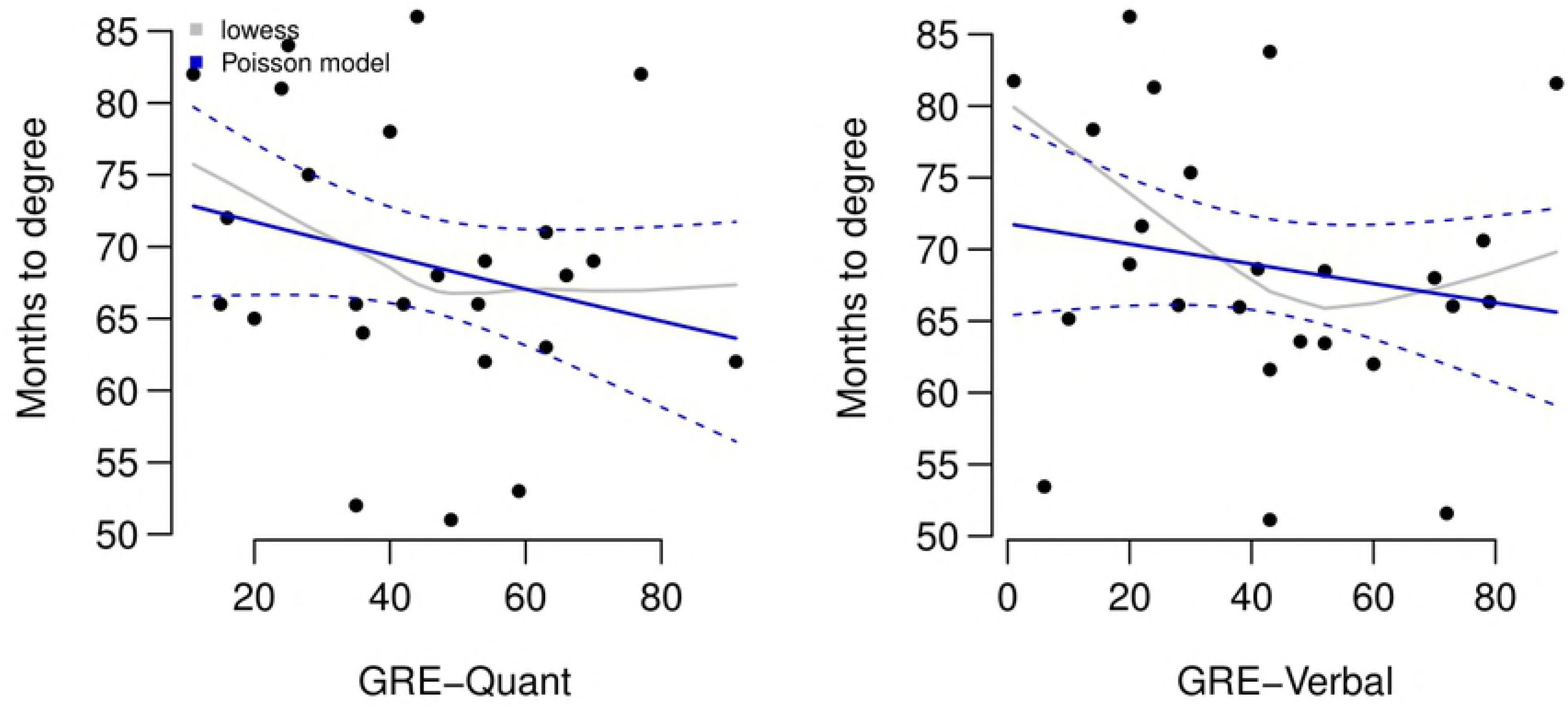
Associations between quantitative and verbal GRE scores and months to degree. Solid curves show the fitted values from the Poisson regression models (dashed lines are 95% confidence intervals) and the grey curves show lowess smoothers (locally weighted scatterplot smoother).

### Lack of association between GRE scores and fellowships

We are well aware that counting papers, either first author or total, has limitations – especially since neither metric captures the quality and/or impact of the publications. Such parameters are difficult to uniformly measure because they are often very field-specific, and sometimes the impact of research is not fully appreciated for years to come. Therefore, we sought to include individual fellowships obtained as one metric of student quality. We included fellowships that are reviewed nationally by panels of experts, providing a comparison between students in this cohort against students at similar stages of training from other institutions around the country. Predoctoral fellowships obtained by this cohort are included in Table 3.

**Table 3:** Individual Fellowships awarded to IMSD students matriculating from 2007-2011.

| **Fellowship type** | **number awarded** |
| --- | --- |
| F31 Ruth L. Kirschstein National Research Service Award (NRSA) Predoctoral Fellowship | 8 |
| American Heart Association Predoctoral Fellowship | 1 |
| National Science Foundation Graduate Research Fellowship | 1 |
| UNCF Merck Graduate Science Research Dissertation Fellowship | 1 |
| Department of Defense Prostate Cancer Research Program Predoctoral Fellowship | 1 |

Boxplots of GRE scores stratified by whether or not students received a fellowship are shown in Fig 6. From bottom to top, the horizontal lines of a boxplot show the min, 25th percentile, median, 75th percentile, and max values in a given group. In Fig 7 the predicted probability of obtaining a fellowship as a function of GRE score is presented. Interestingly, increasing a student’s GRE-Q score by 20 points decreases their odds of receiving a fellowship by 35% (odds ratio = 0.65 with 95% CI 0.27 to 1.58). For instance, the predicted probability of receiving a fellowship for students with GRE-Q scores of 40 and 60 are 50% (95% CI 30% to 71%) and 40% (95% CI is 15% to 64%), respectively. Alternatively, increasing a student’s GRE-V score by 20 points increases their odds of receiving a fellowship by just 4% (odds ratio = 1.04 with 95% CI 0.55 to 1.98). The predicted probability of receiving a fellowship for students with GRE-V scores of 40 vs. 60 are 48% (28%, 68%) and 49% (25%, 73%), respectively. We conclude for this data set, that GRE scores have little value in predicting who will receive a fellowship; in fact, for GRE-Q we observed a negative correlation.

**Fig 6.**
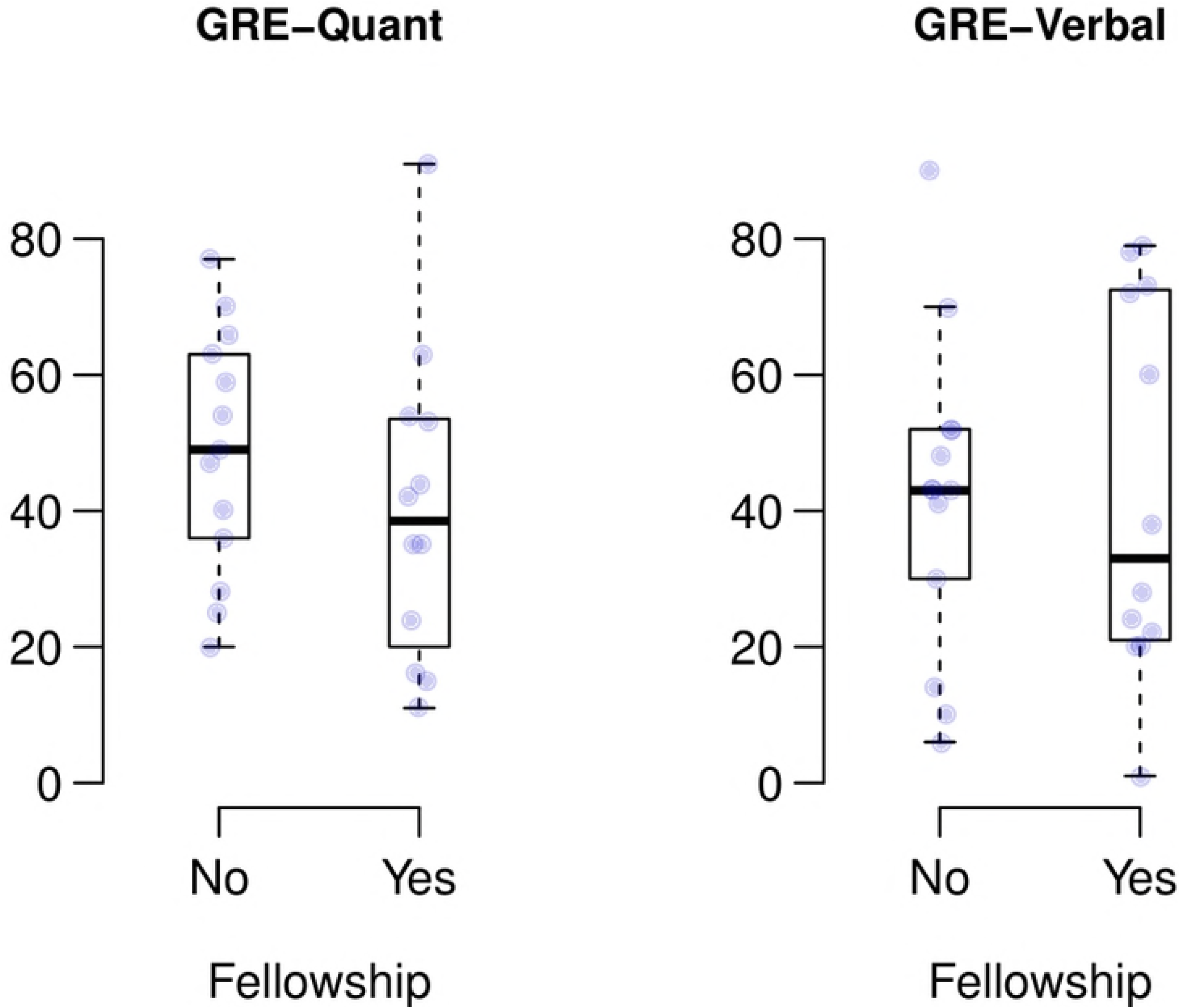
Boxplots of GRE scores stratified by whether or not the students received a fellowship. Data are for fellowships listed in Table 3. The raw data points are overlaid. From bottom to top, the horizontal lines of a boxplot show the minimum GRE score, 25th percentile, median, 75^th^ percentile, and max values in a given group.

**Fig 7.**
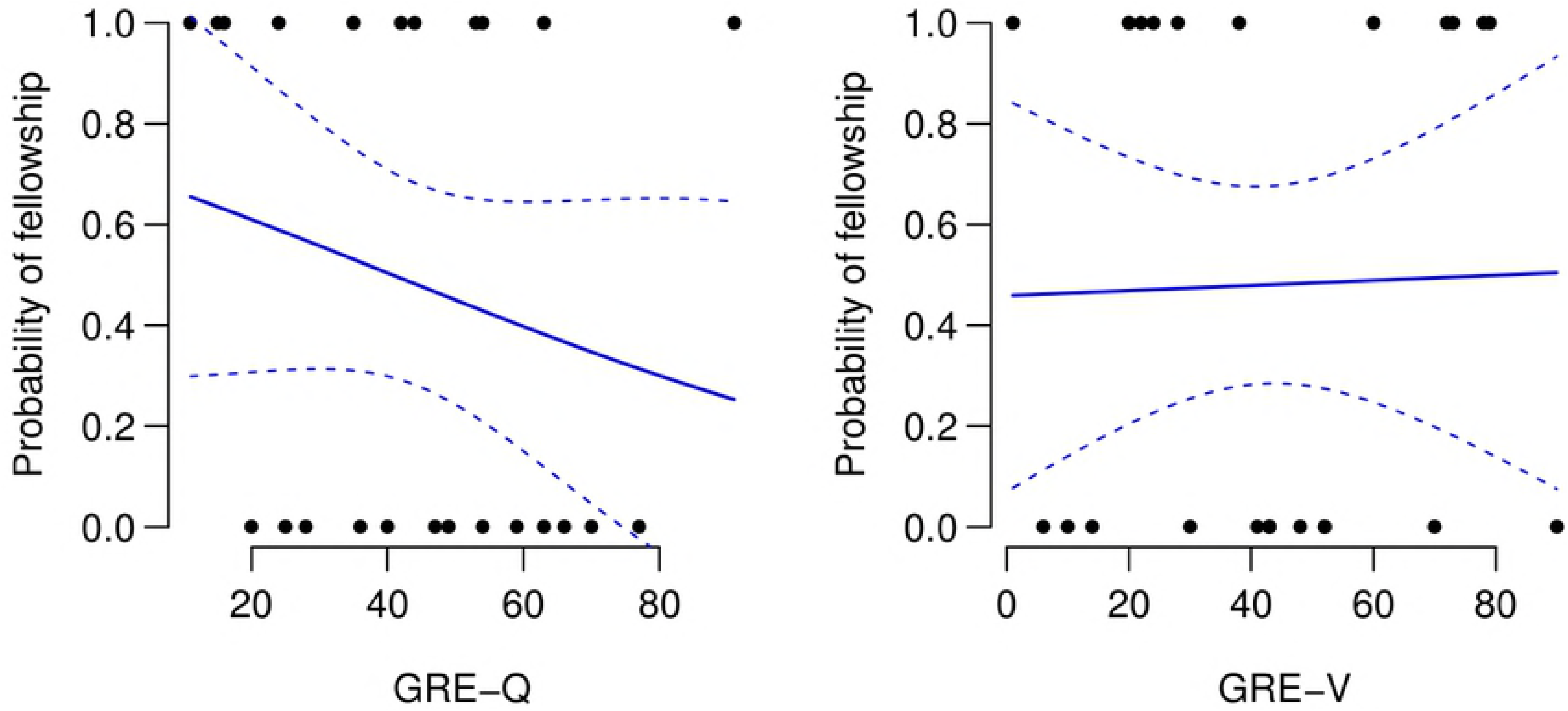
Predicted probability of obtaining a fellowship as a function of GRE scores. Data are for fellowships listed in Table 3. The raw data points are overlaid (0 = No fellowship, 1= Fellowship).

### Lack of association between GRE scores and faculty evaluation

At the completion of their doctoral training, each faculty mentor is asked to evaluate their PhD student on each of ten questions provided in Table 1. The student is not aware that they are or have been evaluated, and the evaluation is never shared with the student nor used for any other purpose. It is important to note that a lower ranking indicates a better evaluation, with 10 being the highest score possible and 50 the lowest score. Fig 8 (left panel) shows the association between GRE-Q score and faculty ranking. As in the prior figures, solid curves show the fitted values from the Poisson regression models (dashed lines are 95% confidence intervals) and the grey lines show the lowess curves. Corresponding data for GRE-V score and faculty ranking are presented in Fig 8, right panel. In each case the associations were small and actually negative (that is, higher GRE scores were associated with lower faculty rankings). Increasing GRE-Q by 20 points increases (worsens) the expected ranking by 5% (rate ratio = 1.05 with 95% CI 0.92 to 1.19). For instance, for students with GRE-Q scores of 40 and 60, the expected rankings are 23.56 and 24.65. Likewise, increasing GRE-V by 20 points increases (worsens) the expected ranking by 10% (rate ratio = 1.10 with 95% CI 1.003 to 1.217). For students with GRE-V scores of 40 and 60, the expected rankings are 23.57 and 26.04.

**Fig 8.**
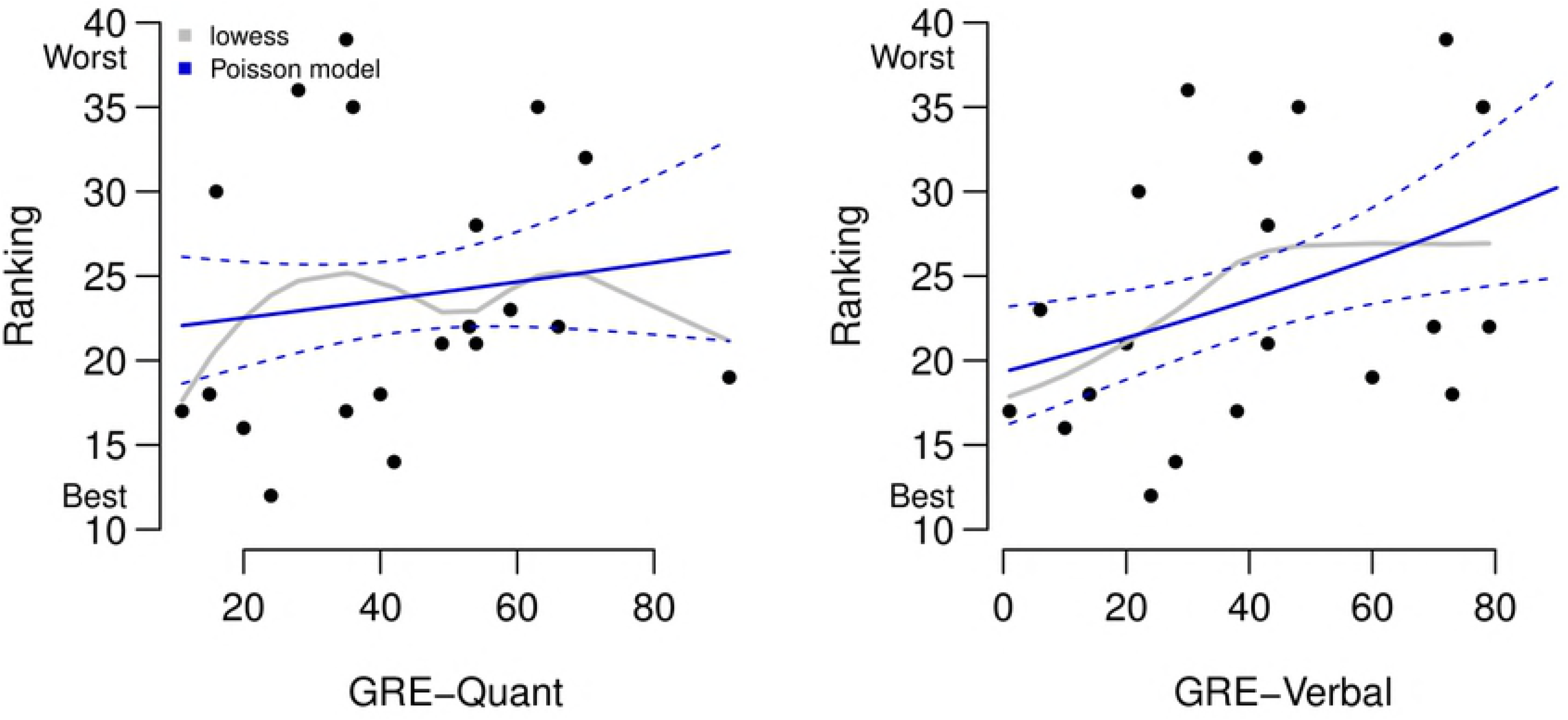
Associations between GRE scores and faculty ranking. Solid curves show the fitted values from the Poisson regression models (dashed lines are 95% confidence intervals) and the grey curves shows lowess smoothers (locally weighted scatterplot smoother).

The data indicate that GRE scores across the entire range of values in this cohort are not predictive of the outcome measures we assessed. We took one final approach – testing for differences in performance measures between the lower and upper quartiles of the GRE scores. To be clear, we compared students with very low scores (<25% GRE-Q or V) to students with very high scores (>75% GRE-Q or V). Although this approach does not use all the data, it would be expected to yield an upwardly biased estimate of the GRE outcome association. The results of such an analysis are shown in Table 4 (for GRE-Q) and Table 5 (for GRE-V). For both tables the first two columns show the mean and standard deviation (SD) of performance measures (number of publications, number of first author publications, months to degree, and faculty ranking) among students in the lower 25th percentile and the upper 25th percentile of GRE score. The third and fourth columns show the difference in mean performance measures between the lower and upper quartiles and the 95% confidence intervals. We see that the point estimates are modest at best, and all confidence intervals include zero as expected. Therefore, even when comparing very low scores, (a range that many graduate schools rarely admit students) to high scores, we do not find evidence that a relationship exists even between the two most likely classes of students.

**Table 4:** Mean (SD) of variables in lower and upper quartiles of GRE-Q and 95% CIs for their difference.

|  | Lower Q | Upper Q | Difference | 95% CI |
| --- | --- | --- | --- | --- |
| No. of publications | 4.6 (2.2) | 4.3 (2.1) | 0.24 | (-2.39, 2.86) |
| No. of first author pubs | 1.7 (0.8) | 1.5 (0.8) | 0.21 | (-0.77, 1.2) |
| Months to degree | 75 (7.6) | 69 (7) | 5.95 | (-2.97, 14.88) |
| Faculty ranking | 21.5 (9.3) | 27 (7.7) | -5.5 | (-18.15, 7.15) |

The first two columns show the mean and standard deviation (SD) of performance measures among students in the lower 25^th^ percentile and the upper 25^th^ percentile. The third and fourth columns show the difference in mean performance measures between the lower and upper quartiles and the 95% confidence intervals.

**Table 5:** Mean (SD) of variables in lower and upper quartiles of GRE-V and 95% CIs for their difference.

|  | Lower Q | Upper Q | Difference | 95% CI |
| --- | --- | --- | --- | --- |
| No. of publications | 5.3 (3.5) | 4.7 (2.3) | 0.62 | (-2.96, 4.2) |
| No. of first author pubs | 2.7 (1.7) | 1.7 (0.8) | 1.05 | (-0.6, 2.69) |
| Months to degree | 72.2 (11.1) | 67.4 (9.6) | 4.86 | (-7.8, 17.52) |
| Faculty ranking | 20.8 (5.2) | 27.2 (9.2) | -6.37 | (-17.67, 4.93) |

The first two columns show the mean and standard deviation (SD) of performance measures among students in the lower 25^th^ percentile and the upper 25^th^ percentile. The third and fourth columns show the difference in mean performance measures between the lower and upper quartiles and the 95% confidence intervals.

### Outcomes of the cohort to date

For the 25 students in the cohort analyzed here, the final question we can ask is where are they now? As mentioned earlier, most of the cohort moved on to a postdoctoral position upon PhD completion at a range of research intensive institutions listed in Table 6. The students in this cohort completed their PhDs between spring 2012 and summer 2017, so some have had time to move to a position beyond the first postdoc. So far after completing their first postdoctoral position, two individuals have moved on to Biopharma, one who is developing a start-up company, one moved to an administrative position at NIH, and one is now a tenure-track assistant professor. At of the time of this writing (June 2018), none of this cohort of 25 students have left science.

**Table 6:** Postdoctoral institutions for IMSD students upon PhD completion.

|  |
| --- |
| Harvard University |
| Mt Sinai Icahn School of Medicine |
| University of Texas Health Science Center |
| Northwestern University |
| Yale University |
| University of Florida |
| Vanderbilt University Medical Center |
| John Hopkins University |
| National Institutes of Health |
| Baylor College of Medicine |
| University of Colorado |
| Michigan State University |
| St Jude Children’s Research Hospital |
| Case Western University |
| University of Washington |
| Charles R. Drew University of Medicine and Science |
| Vanderbilt University |

Institutions where IMSD students who matriculated from 2007-2011 completed first postdocs

## Discussion

As a result of the admissions process adopted by the Vanderbilt IMSD program over a decade ago, we now have a cohort of graduate students whose GRE scores spanned the entire range from 1-91 percentile who have completed the PhD. This analysis includes 25 IMSD students who matriculated into our biomedical research programs from 2007-2011 and completed PhDs beginning in 2012 to summer 2017. We consistently observed only associations between academic outcomes and GRE scores. Even when accounting for the variability in these estimates (i.e., the width of the 95% CI) we see that the data support only very minor associations, if any. This can be visualized by looking at the confidence bands for the regression lines. For example, when modeling the number of first author publications as a function of quantitative GRE score, we found the rate ratio (slope) was 1.004 (95% CI 0.994 to 1.014). This implies that the average change in the number of first author publications is nearly zero even for a large shift in the GRE quantitative percentile. However, the data support changes of approximately [-1 to +1] publication. While not exactly zero, these limited data clearly support the hypothesis that there is only a very minor relationship, if any, between publication and GRE scores. In fact, for verbal scores we observed a very small negative relationship indicating (not statistically significant from zero) that there is essentially no association in these data. Similar findings can be observed for the other outcome metrics presented here, including first author papers, fellowships obtained, time to degree, and faculty evaluations at exit. Importantly, we did observe a statistically significant relationship in the opposite direction with GRE and ranking (better ranked individuals tended to have poorer GRE scores). So while the overall sample size is small, there appears to be enough precision or power in these data to detect strong associations if they existed.

We have evaluated verbal and quantitative GRE scores separately in this study, but in actuality a student’s application contains both scores. Perhaps a very low score in one domain (Q or V) may be offset by a high score in the other. In fact, most of the students in this cohort had two reasonably comparable Q and V scores. Of the 25 students, only four had a percentile spread between their two scores of greater than 30. In other words, they were generally either poor test takers or strong ones. Furthermore, only six of the students who completed PhDs had both GRE-Q and GRE-V scores above the 50^th^ percentile, making it questionable whether the other 19 would have gained admittance to a graduate program that adhered to higher expectations for GRE performance. Five of the 25 students had neither GRE-Q or GRE-V scores above the 30^th^ percentile. We think it unlikely that they would be offered admission to most graduate programs at the time or even to many programs today. Yet, among this group of five is the student who garnered the best (lowest score) faculty evaluation. These outcomes underscore the benefit of giving letters of recommendation, personal statements, and interviews far more weight than GRE scores in making admissions decisions. Our GRE-tolerant approach for increasing the number of students from historically underrepresented groups completing PhDs has been highly successful.

The relationship between objective test scores and performance has been a subject of debate for many years. Uncertainty surrounding their predictive ability must be weighed against the cost imposed on applicants to take the test, and the advantages available to a subset of applicants who can prepare extensively ahead of time and/or take the test multiple times to obtain the desired high scores. However, the outcomes of the cohort presented here indicate that non-quantitative measures (letters of recommendation, personal statements, interviews) are capable of selecting successful PhD candidates, even when those candidates have extremely low GRE scores. Subjective measures have their own drawbacks, and we sought to minimize these by having multiple, experienced readers of graduate student applications. We attempted to mediate individual biases by including multiple diverse viewpoints of each student’s potential in reaching a decision to offer admission. Admittedly, this process is time consuming, but the decision of who to train as the next generation of PhD scientists is also arguably one of the most important we make.

The “GRExit” movement is growing, and for those biomedical programs that remain undecided, the data here may be helpful in arriving at a decision on whether or not to continue to require GRE scores for admission. However these decisions turn out, we assert that our GRE-tolerant approach (no score too low) undoubtedly opened doors of opportunity for PhD training at Vanderbilt that may have otherwise remained closed for historically underrepresented students with very low GRE scores. The increased diversity they bring to the community of PhD biomedical scientists will be a benefit for decades to come.

## Acknowledgments

We thank Richard Hoover, Associate Dean of the Graduate School, for his unwavering support of the IMSD program and for working with the IMSD admissions committee to matriculate IMSD students into the Graduate School. We also thank Cathleen Williams, IMSD program manager during 2007-11, for her invaluable help and support. Finally we thank the supportive faculty mentors and the talented IMSD students from this period for their hard work and achievements that have been reported in this study. This study was supported by NIH R25GM062459 to RC and LS.

## Supporting information

**Table S1.** (corresponds to Fig 3. Associations between quantitative and verbal GRE scores and total number of publications)

Results from Poisson regression models looking at the association between GRE-Quantitative and number of publications (Table 1a) and GRE-Verbal and number of publications (Table 1b). The columns show the estimated rate ratios, model robust standard errors, 95% confidence intervals, and p-values.

**Table S2.** (corresponds to Fig 4. Associations between quantitative and verbal GRE scores and total number of first author publications)

Results from Poisson regression models looking at the association between GRE-Quantitative and number of first author publications (Table 2a) and GRE-Verbal and number of first author publications (Table 2b). The columns show the estimated rate ratios, model robust standard errors, 95% confidence intervals, and p-values.

**Table S3.** (corresponds to Fig 5. Associations between quantitative and verbal GRE scores and months to degree)

Results from Poisson regression models looking at the association between GRE-Quantitative and number of months to degree (Table 3a) and GRE-Verbal and months to degree (Table 3b). The columns show the estimated rate ratios, model robust standard errors, 95% confidence intervals, and p-values.

**Table S4.** (corresponds to Fig 6. Boxplots of GRE scores stratified by whether or not the students received a fellowship)

Results from logistic regression models looking at the association between GRE-Quantitative and receipt of fellowship (Table 4a) and GRE-Verbal and receipt of fellowship (Table 4b). The columns show the estimated odds ratios, model robust standard errors, 95% confidence intervals, and p-values.

**Table S5.** (corresponds to Fig 8. Associations between GRE scores and faculty ranking) Results from Poisson regression models looking at the association between GRE-Quantitative and faculty ranking (Table 5a) and GRE-Verbal and faculty ranking (Table 5b). The columns show the estimated rate ratios, model robust standard errors, 95% confidence intervals, and p-values.

